# Analysis of motor unit spike trains estimated from high-density surface electromyography is highly reliable across operators

**DOI:** 10.1101/2021.02.19.431376

**Authors:** François Hug, Simon Avrillon, Alessandro Del Vecchio, Andrea Casolo, Jaime Ibanez, Stefano Nuccio, Julien Rossato, Aleš Holobar, Dario Farina

## Abstract

There is a growing interest in decomposing high-density surface electromyography (HDsEMG) into motor unit spike trains to improve knowledge on the neural control of muscle contraction. However, the reliability of decomposition approaches is sometimes questioned, especially because they require manual editing of the outputs. We aimed to assess the inter-operator reliability of the identification of motor unit spike trains. Eight operators with varying experience in HDsEMG decomposition were provided with the same data extracted using the convolutive kernel compensation method. They were asked to manually edit them following established procedures. Data included signals from three lower leg muscles and different submaximal intensities. After manual analysis, 126 ± 5 motor units were retained (range across operators: 119-134). A total of 3380 rate of agreement values were calculated (28 pairwise comparisons × 11 contractions/muscles × 4-28 motor units). The median rate of agreement value was 99.6%. Inter-operator reliability was excellent for both mean discharge rate and time at recruitment (intraclass correlation coefficient > 0.99). These results show that when provided with the same decomposed data and the same basic instructions, operators converge toward almost identical results. Our data have been made available so that they can be used for training new operators.

## Introduction

Movements result from the activation of motor units, each comprising of a motoneuron and the muscle fibers it innervates. Motoneurons send trains of action potentials to the muscle fibers, which produce force. The neural drive is the net output of all the motor neurons that innervate the muscle, i.e. the ensemble of activation times of the motoneurons. In healthy individuals, there is a one-to-one relationship between the generation of an action potential in the motor neuron and the generation of an action potential in the innervated muscle fibers. Thus, the discharge times of motor units contain direct information about the neural drive. As a result, there is a growing interest in decoding the discharge characteristics of motor units to advance our knowledge on the neural control of movement (Del Vecchio et al., 2020; Farina et al., 2016), to develop human-machine interfaces (Chen et al., 2020), and to design rehabilitation interventions in people with neurological conditions (Ting et al., 2019).

Activity of motor units is conventionally detected using intramuscular electromyography (EMG) electrodes. Although it provides direct recordings of motor unit action potentials, this invasive technique can only identify a limited number of units from small muscle regions. Recent developments in electrode technology and signal processing makes it possible to identify the concurrent activity of many motor units non-invasively (Farina et al., 2016; Marateb et al., 2011). Specifically, high-density surface EMG (HDsEMG) electrodes can be used to provide a spatial sampling of motor unit action potentials (Merletti et al., 1999; Merletti et al., 2008). Blind source separation procedures (Holobar and Zazula, 2007; Negro et al., 2016) applied to these signals currently enable the identification of up to 30-40 motor units. However, after an automatic extraction, most of these approaches require some degree of manual analysis/editing of the motor unit spike trains to check for false positives and false negatives (Del Vecchio et al., 2020; Enoka, 2019). This manual analysis consists of reinforcing the motor unit spike trains with tuned motor unit separation filters. The motor unit separation filter is a spatio-temporal filter that, when applied to surface EMG, yields the spike train of individual motor unit. The coefficients of the motor unit filters form the inverse of the surface EMG mixing matrix and are estimated automatically by decomposition algorithms. Afterwards, each spike train is segmented into motor unit firing patterns and baseline noise. Each individual motor unit filter can then be recalculated from the segmented firings of a given motor unit. In this process, some of the automatically segmented motor unit firings may be manually removed or additional ones added, refining the motor unit filter. The refined motor unit filter may be reapplied to surface EMG signals, yielding refined motor unit spike train and its segmentation into motor unit firings. This procedure is iteratively repeated, leading to manual improvement of motor unit firing patterns. In motor unit filter refinement, manual adding or removing of the motor unit firings is typically based on the height of the spikes in the identified spike train and on pulse-to-noise ratio (Holobar et al., 2014), but may also consider regularity of inter-spike interval of identified motor unit firing patterns. This manual step potentially calls into question the accuracy of such results, casting doubt on any conclusions reached by interpreting decomposed motor unit activity. It is therefore important to quantify the reliability of the manual editing step among different operators. In other words, we need to determine whether different operators who have access to the same output from the decomposition algorithm would extract similar motor unit spike trains.

The overall aim of this study was to assess the inter-operator reliability of the identification of motor unit spike trains. Specifically, we aimed to assess the rate of agreement among operators from experimental signals collected during submaximal contractions and its relationship with the accuracy of the motor unit identification (pulse-to-noise ratio). A secondary aim was to assess the inter-operator reliability for two motor unit discharge characteristics commonly used in the literature, i.e., the mean discharge rate and the time at which the first action potential is detected. All raw and processed data have been made available so that they can be used for training new operators.

## Methods

### 1. Participants

Eight operators with varying amount of experience in HDsEMG decomposition volunteered to participate in this experiment (Table 1). They all co-authored this article (FH, JI, JR, SN, SA, ADV, AC, and AH). The research ethics committee of the University of Nantes approved this study (CERNI, n°13102020), which was based on the re-use of data collected in two different experiments (approval # 2013001448 from the University of Queensland and approval #44 680 from the University of Rome ‘Foro Italico’).

**Table 1.** Operator experience. The number of (co)authored articles that involved decomposition of HDsEMG signals is given for each operator, together with the total time each operator took to edit the motor units in the present study. Note that each operator was previously exposed to manual analysis of motor unit spike trains, with varying level of experience as reflected in their number of (co)authored articles.

| Operator # | Background | No. studies with HDsEMG decomposition | Time to edit (min) |
| --- | --- | --- | --- |
| 1 | Human movement science | 2 | 239 |
| 2 | Informatics and systems engineering | 2 | 310 |
| 3 | Human movement science | 0 | 210 |
| 4 | Human movement science | 0 | 389 |
| 5 | Human movement science | 2 | 428 |
| 6 | Human movement science | 20 | 186 |
| 7 | Human movement science | 5 | 211 |
| 8 | Computer science | 55 | 152 |

### 2. Experimental data

In two different experiments, HDsEMG was recorded from the Gastrocnemius lateralis (GL) and Gastrocnemius medialis (GM) [experiment I, (Hug et al., 2021)] and the tibialis anterior (TA) [experiment II, (Del Vecchio et al., 2019)]. These muscles were selected because they usually yield to different numbers – and accuracy - of decomposed units. For example, Hug et al. (2021) highlighted the challenge of extracting motor units from the GL muscle. In contrast, the TA muscle is known to be a reliable muscle for HDsEMG decomposition (Del Vecchio et al., 2020). Apart from the muscle investigated, the decomposition output depends on the contraction level and properties of the subcutaneous layers, which are specific to each participant (Del Vecchio et al., 2020). Therefore, we focused on data from four contraction intensities collected from different participants. Of note, the two original studies (Del Vecchio et al., 2019; Hug et al., 2021) measured only males and therefore the present study only considers decomposition from data collected in males.

#### 2.1. Experimental design

Data from the GL and GM muscles were collected during a series of isometric tasks, as detailed in Hug et al. (2021). Participants laid prone on a custom-made dynamometer equipped with a torque sensor (TRE-50K, Dacell, Korea). Their right ankle angle was set at 10° of plantarflexion (0° being the foot perpendicular to the shank), with their knee was fully extended. After a warm-up involving 15-20 submaximal contractions, the participants performed three maximal isometric contractions for 3 to 5 s with 120-s rest in between. The maximal value obtained from a moving-time window of 250-ms was considered as the peak torque (MVC torque). Then, participants performed three contractions at each of the following intensities: 10%, 30%, 50%, and 70% of their MVC torque. The order of the intensities was randomized. These contractions involved a 5-s ramp-up, a 15-s (50% and 70% of MVC) or 20-s plateau (10% and 30% of MVC) and a 5-s ramp down phase. The contractions were separated by either 60-s (10% of MVC) or 120-s (30%, 50% and 70% of MVC) of rest. Feedback of the target and torque output was displayed on a monitor.

The detailed description of the experimental protocol with the TA muscle has been described previously (Del Vecchio et al., 2019). The force sensor (load cell, CCt Transducer s.a.s, Turin, Italy) was placed in a custom dynamometer to measure the isometric dorsiflexion force of the dominant leg [see Figure 1 in (Del Vecchio et al., 2019)]. Participants were seated on a massage table with their hip flexed at 120° (180° being supine), their knee extended to 180°, and their ankle placed at 10° of plantar flexion (0° being the foot perpendicular to the shank). The foot and the ankle were held in place with Velcro straps on an adjustable footplate. Participants performed three to four maximal isometric contractions with 30 s of recovery in between. The peak force (MVC force) was used as a reference to determine the target force for the submaximal contractions. These contractions involved a ramp-up phase at 5% MVC s^−1^, a 10-s plateau (35%, 50%, or 70% MVC), and a ramp-down phase performed at the same rate as the ramp-up phase. Two repetitions were performed at each contraction level. The order was randomized, and the contractions were separated by 3-5 min of rest.

**Fig 1.**
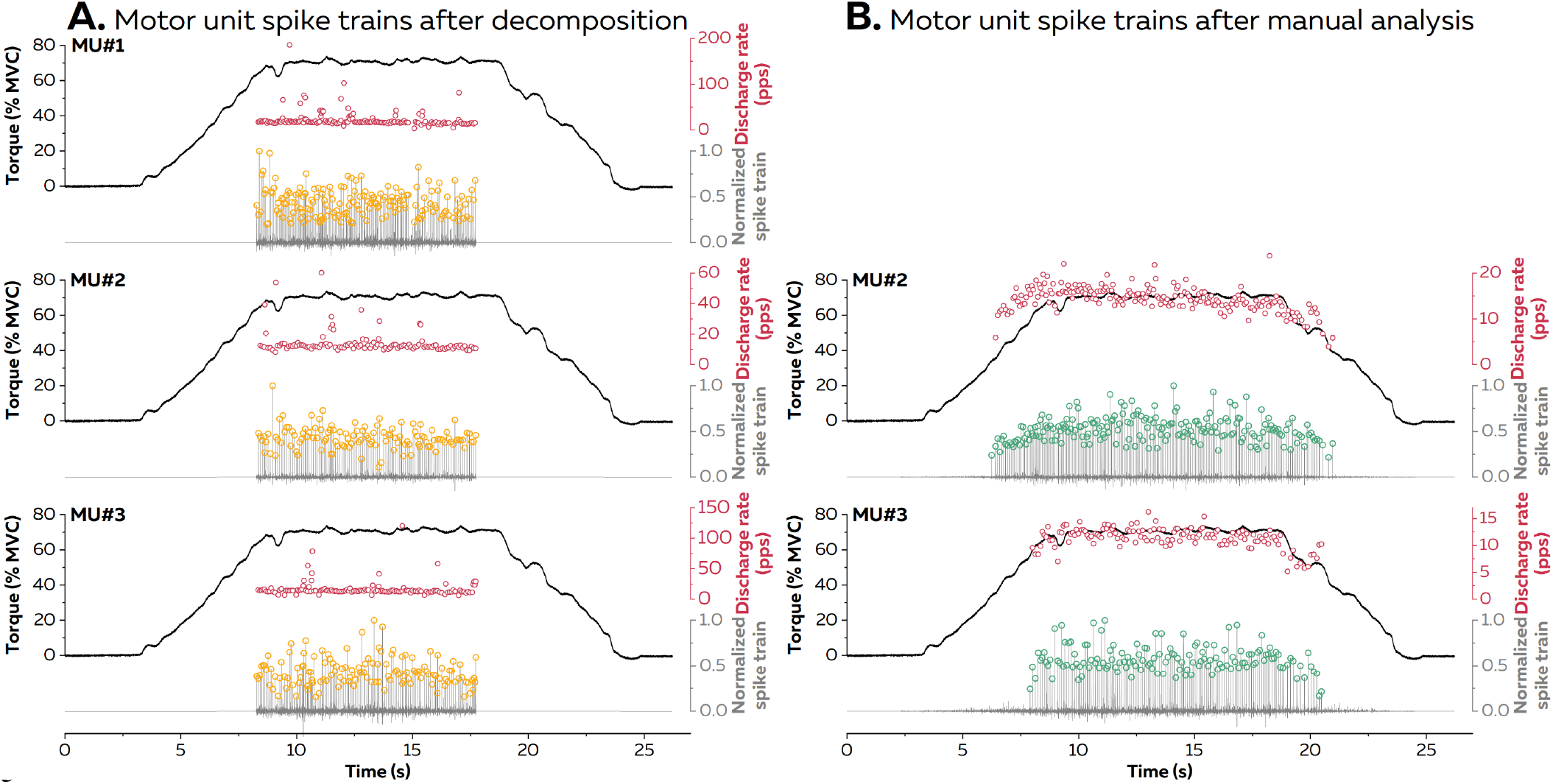
Example of the decomposition outcome before and after manual analysis. Three motor units from the gastrocnemius medialis (70% MVC) are depicted. The left panels (A) exhibit the discharge times identified by the automatic decomposition over a 10-s window. After identifying and removing the spikes with lower quality, the motor-unit filter was re-calculated and re-applied over the whole contraction. The new spikes recognized as motor unit firings were added leading to the results depicted on the right panels (B). Of note, manual analysis of the first motor unit (MU#1) did not lead to a pulse-to-noise ratio>30 dB, and therefore this motor unit was discarded following manual editing. MVC, Maximal Voluntary Contraction.

#### 2.2. High-density surface electromyography

For GL, GM, and TA muscles, HDsEMG was measured using two-dimensional adhesive grids of 64 electrodes (13×5 electrodes with one electrode absent from a corner, gold-coated, inter-electrode distance: 8 mm; [ELSCH064NM2, SpesMedica, Battipaglia, Italy]).

The grids were aligned with the main fascicle direction as determined using B-mode ultrasound (Aixplorer, Supersonic Imagine, France) for GM and GL or using a dry array of 16 electrodes for TA. Before electrode application, the skin was shaved and then cleaned with an abrasive pad and alcohol. The adhesive grids were held on the skin using semi-disposable bi-adhesive foam layers (SpesMedica, Battipaglia, Italy). The skin-electrode contact was optimized by filling the cavities of the adhesive layers with conductive paste (SpesMedica, Battipaglia, Italy). Strap electrodes dampened with water were placed around the contralateral (ground electrode) and ipsilateral ankle (reference electrode). The EMG signals were recorded in monopolar mode, bandpass filtered (20-500 Hz) and digitized at a sampling rate of 2048 Hz using a multichannel acquisition system (EMG-Quattrocento; 400-channel EMG amplifier, OT Biolelettronica, Turin, Italy).

## 3. Data analysis

### Automatic decomposition of the HDsEMG signals

First, the monopolar EMG signals were bandpass filtered between 20 and 500 Hz using a second-order Butterworth filter. The HDsEMG signals were decomposed with the convolutive blind source separation method (Holobar and Farina, 2014; Holobar et al., 2014; Holobar and Zazula, 2007) implemented in the DEMUSE software tool (v5.01; The University of Maribor, Slovenia). The following extraction parameters were selected: 50 iterations, maximal coefficient of variation of 50%. This decomposition procedure can identify motor unit discharge times over a wide range of contraction intensities and has been extensively validated using experimental and simulated signals (Holobar and Farina, 2014; Holobar et al., 2014). The decomposition was performed on either 10-s (TA, GL, and GM at 50% and 70% MVC) or 15-s (GL and GM at 10% and 30% MVC) centered on the torque (GL, GM) or force (TA) plateau.

### Manual edition of the decomposition results

After the automatic identification of the motor units, all the motor unit spike trains were visually inspected and manually edited by each of the eight operators. Together with the initial automatic decomposition data, a standardized list of instructions was given to each operator. Specifically, they had to:

i. Read a tutorial (Del Vecchio et al., 2020) to make sure that they all had the same basic knowledge on HDsEMG decomposition,
ii. Inspect and edit the motor unit spike trains over the whole contraction,
iii. Remove unreliable motor units that had a pulse-to-noise ratio lower than 30 dB (Holobar et al., 2014), as classically done in the literature (Avrillon et al., 2021; Laine et al., 2015),
iv. Note the time that they took to edit each file.

It should be noted that the decision to discard a motor unit with a pulse-to-noise ratio < 30 was taken after manual edition (step ii), i.e., after first trying to improve its accuracy. As explained in detail in Del Vecchio et al. (2020), the manual editing/analysis of the motor unit spike trains (step ii) consisted in the following steps performed in an iterative way: i) identifying and removing the spikes of lower quality, ii) re-calculating the motor-unit filter and re-applying it over a portion of the signal, and iii) adding the new spikes recognized as motor unit firings. Each of these steps includes subjective decision-making, which is a potential source of discrepancies between operators.

### Rate of agreement

For each motor unit, the rate of agreement (RoA) between decompositions was calculated for all pairs of operators (i.e. 28) as follows:

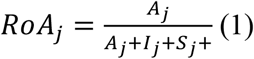

Where *Aj* denotes the number of discharges of the *j*th motor unit that were identified by both operators, *Ij* is the number of discharges identified by operator #1 but not by operator #2, and *Sj* is the number of discharges identified by operator #2 but not by operator #1. The discharge time tolerance was set to ±1 time point. It means that a firing was considered *Aj* if it was identified by both operators at time instants separated by not more than ∼ 0.49 ms (1/2048 Hz). Of note, a total of 3380 RoA values were calculated, i.e. 28 pairwise comparisons × 11 contractions × 4-28 motor units per contraction.

### Motor unit discharge characteristics

We assessed the inter-operator reliability of two motor unit discharge characteristics commonly extracted from motor unit spike trains. First, the time of recruitment of each motor unit was determined as the time when the first action potential was observed. Second, we estimated the mean discharge rate of each motor unit during the plateau of torque/force.

## 5. Statistical analysis

Statistical analyses were performed in Statistica v7.0 (Statsoft, Tulsa, OK, USA). A Shapiro-Wilk test was used to test for a normal distribution. A Mann-Whitney test was used to compare the RoA and the pulse-to-noise ratio values between the motor units that had been identified by all eight operators and those that had been only identified by some of the operators. We used Pearson’s correlation coefficient to assess the relationship between the RoA and the pulse-to-noise ratio of the edited units. Finally, we assessed the inter-operator reliability of the motor unit discharge rate and time of motor unit recruitment by calculating the intraclass coefficient of correlation (ICC). All data are reported as mean ± standard deviation and the level of significance was set at p ≤0.05.

## Results

The entire dataset is available at <u>10.6084/m9.figshare.13695937</u>

Fig. 1 depicts an example of the outcome of the decomposition, before and after the manual analysis step. The automatic decomposition allowed the identification of 231 motor units (ranging from 5 for GL at 10% MVC to 39 for GM at 30% MVC; Table 2). On average, 265 ± 99 min were needed for manual editing of these motor units (Table 1), i.e., ≈1 min 8s per motor unit for contractions ranging from 25 to 30 s. After manual editing, 126 ± 5 motor units were retained for further analysis (range across operators: 119-134) (Table 2). The number of motor units identified by each operator is detailed in Table 3.

**Table 2.** Characteristics of the decomposed motor units. The number of decomposed motor units (No. MUs), the mean pulse-to-noise ratio (PNR), and the mean discharge rate (DR) obtained both before and after the manual editing are given for each contraction. The mean discharge rate was calculated over the force/torque plateau. Of note, the mean discharge rate after automatic decomposition is reported only for the motor units which were retained after manual editing and all instantaneous discharge rate values > 100pps were automatically removed. GL, Gastrocnemius lateralis; GM, Gastrocnemius medialis; TA, Tibialis anterior; MUs, motor units. MVC, maximal voluntary contraction

| Muscle & condition | After automatic decomposition |  |  | After manual editing |  |  |
| --- | --- | --- | --- | --- | --- | --- |
|  | No. MUs | Mean PNR (range) | Mean DR, pps (range) | No. MUs (range) | Mean PNR (range) | Mean DR, pps (range) |
| **GL 10% MVC** | 5 | 34.2±5.1<br>(27.1-39.8) | 10.1±1.9<br>(8.6-12.5) | 4.0±0<br>(4-4) | 37.5±7.5<br>(31.2-47.2) | 8.1±0.9<br>(7.2-8.9) |
| **GL 30% MVC** | 16 | 30.7±5.2<br>(18.9-37.7) | 10.8±1.6<br>(8.5-12.8) | 8.6±0.7<br>(8-10) | 33.7±2.2<br>(30.7-36.8) | 8.2±1.3<br>(5.2-9.6) |
| **GL 50% MVC** | 36 | 23.5±5.9<br>(16.2-35.2) | 15.7±2.7<br>(13.4-19.5) | 4.0±0<br>(4-4) | 32.5±1.8<br>(30.5-34.7) | 11.3±1.3<br>(10.1-12.8) |
| **GL 70% MVC** | 13 | 29.8±4.5<br>(20.5-36.3) | 17.9±1.1<br>(16.8-19.4) | 4.9±0.4<br>(4-5) | 32.5±1.1<br>(31.7-34.3) | 14.0±1.5<br>(12.5-15.7) |
| **GM 10% MVC** | 22 | 31.9±3.1<br>(26.4-38.6) | 10.8±1.5<br>(8.7-13.2) | 16.5±0.8<br>(16-18) | 32.6±2.0<br>(30.4-37.5) | 8.1±1.1<br>(6.3-10.4) |
| **GM 30% MVC** | 39 | 31.8±6.1<br>(18.9-41.2) | 11.9±2.2<br>(7.7-17.3) | 26.1±1.6<br>(24-28) | 34.6±2.7<br>(30.0-40.3) | 8.1±1.0<br>(6.0-10.5) |
| **GM 50% MVC** | 19 | 37.4±6.4<br>(23.3-46.5) | 11.7±3.2<br>(7.2-18.6) | 15.4±0.5<br>(15-16) | 38.1±4.3<br>(31.2-45.1) | 7.6±1.2<br>(5.7-9.8) |
| **GM 70% MVC** | 10 | 31.3±3.5<br>(27.3-39.2) | 16.5±1.6<br>(14.2-18.4) | 5.8±0.5<br>(5-6) | 32.4±2.8<br>(30.2-37.7) | 12.2±0.6<br>(11.7-13.1) |
| **TA 35% MVC** | 22 | 37.8±4.8<br>(27.8-49.6) | 15.6±3.0<br>(9.0-21.5) | 20.3±1.0<br>(19-21) | 38.7±4.0<br>(32.8-50.1) | 12.0±1.2<br>(9.0-14.4) |
| **TA 50% MVC** | 18 | 32.6±5.4<br>(18.8-39.0) | 18.6±2.5<br>(13.7-23.3) | 13.3±1.3<br>(12-15) | 35.6±3.4<br>(31.0-40.6) | 15.5±1.4<br>(12.4-17.6) |
| **TA 75% MVC** | 31 | 24.3±6.8<br>(17.3-36.9) | 24.5±4.6<br>(14.2-29.6) | 6.9±0.8<br>(6-8) | 33.4±1.9<br>(30.7-36.1) | 17.4±1.5<br>(15.4-20.5) |

**Table 3.** Number of selected motor units per operator. The operators were asked to select motor units only if their pulse-to-noise ratio was higher than 30 (see Methods). Despite this common criterion, the manual decomposition yielded to a slightly different number of units depending on the operators. MUs, motor units. MVC, maximal voluntary contraction

|  | Operators |  |  |  |  |  |  |  |
| --- | --- | --- | --- | --- | --- | --- | --- | --- |
|  | #1 | #2 | #3 | #4 | #5 | #6 | #7 | #8 |
| **GL 10% MVC** | 4 | 4 | 4 | 4 | 4 | 4 | 4 | 4 |
| **GL 30% MVC** | 9 | 10 | 9 | 8 | 8 | 8 | 8 | 9 |
| **GL 50% MVC** | 4 | 4 | 4 | 4 | 4 | 4 | 4 | 4 |
| **GL 70% MVC** | 5 | 5 | 5 | 4 | 5 | 5 | 5 | 5 |
| **GM 10% MVC** | 16 | 17 | 16 | 16 | 18 | 16 | 16 | 17 |
| **GM 30% MVC** | 27 | 28 | 27 | 25 | 27 | 24 | 24 | 27 |
| **GM 50% MVC** | 16 | 16 | 15 | 15 | 16 | 15 | 15 | 15 |
| **GM 70% MVC** | 6 | 6 | 5 | 5 | 6 | 6 | 6 | 6 |
| **TA 35% MVC** | 21 | 21 | 21 | 19 | 21 | 19 | 19 | 21 |
| **TA 50% MVC** | 14 | 15 | 12 | 12 | 12 | 13 | 13 | 15 |
| **TA 70% MVC** | 8 | 8 | 6 | 7 | 6 | 6 | 7 | 7 |
| **Total number of MUs** | **130** | **134** | **124** | **119** | **127** | **120** | **121** | **130** |

Fig. 2 shows representative motor unit spike trains identified by each of the eight operators. A total of 3380 RoA values were calculated (see Methods). The mean and median RoA values were 98.9% and 99.6%, respectively (Fig 3). RoA ranged from 56.5% to 100%, with only 20 out of 3380 values (0.6%) being lower than 85%. These lowest RoA values were observed for two particular motor units (motor unit #4 for GL 50% MVC and motor unit #15 for GM 50% MVC). For motor unit #15, one operator did not recalculate the motor unit filter over the whole contraction. This led to some firings being missed, and thus to a low RoA value (60.2%) for the comparison with each of the seven other operators. Of note, the RoA was 100% when calculated between all the other operators. For motor unit #4, the recalculation of the motor unit filter led to the identification of new spikes with moderate quality, which were selected by two out of the eight operators (Fig. 4).

**Fig 2.**
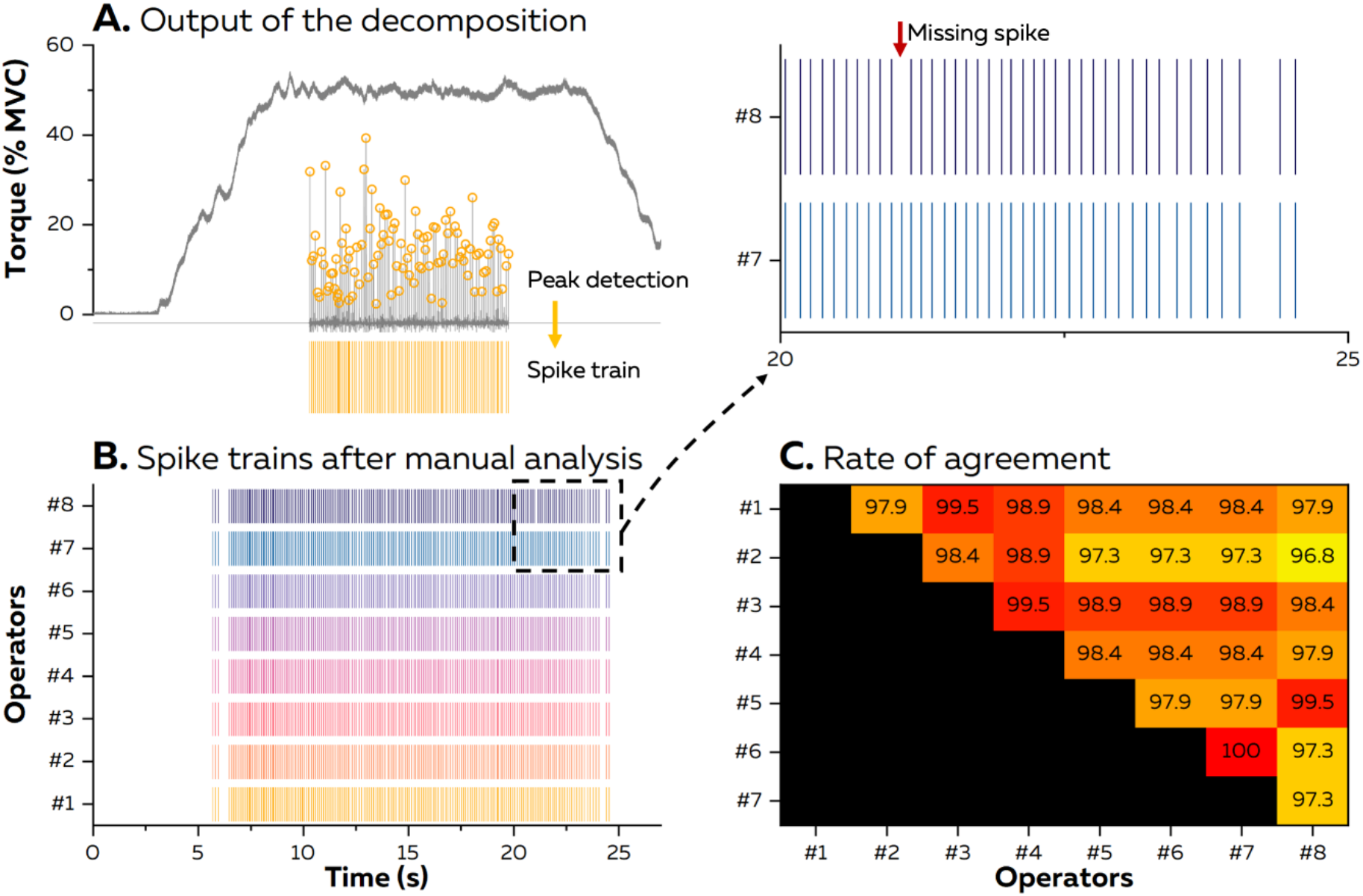
Example of the inter-operator reliability of the identification of a motor unit spike train (MU #2 of the Gastrocnemius medialis). After the automatic identification of the motor units on a portion of the torque plateau (Panel A), all the motor unit spike trains were visually inspected and manually edited by each of the eight operators (Panel B). The rate of agreement was then calculated for each pair of operators (Panel C).

**Fig 3.**
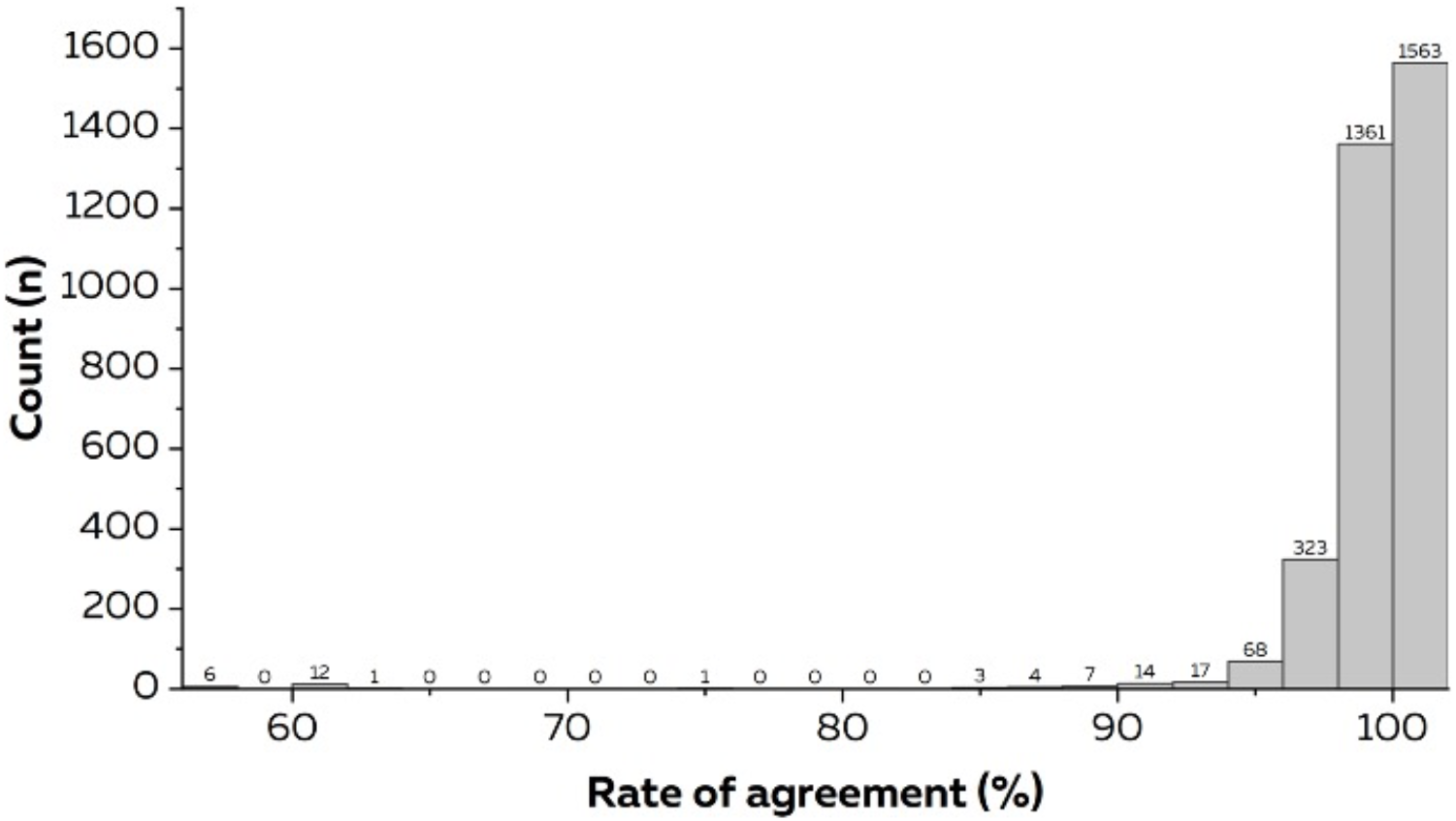
Distribution of rate of agreement values. The rate of agreement was calculated for each pair of operators leading to 3380 values (28 pairwise comparisons × 11 contractions × 4-28 motor units per contraction).

**Fig 4.**
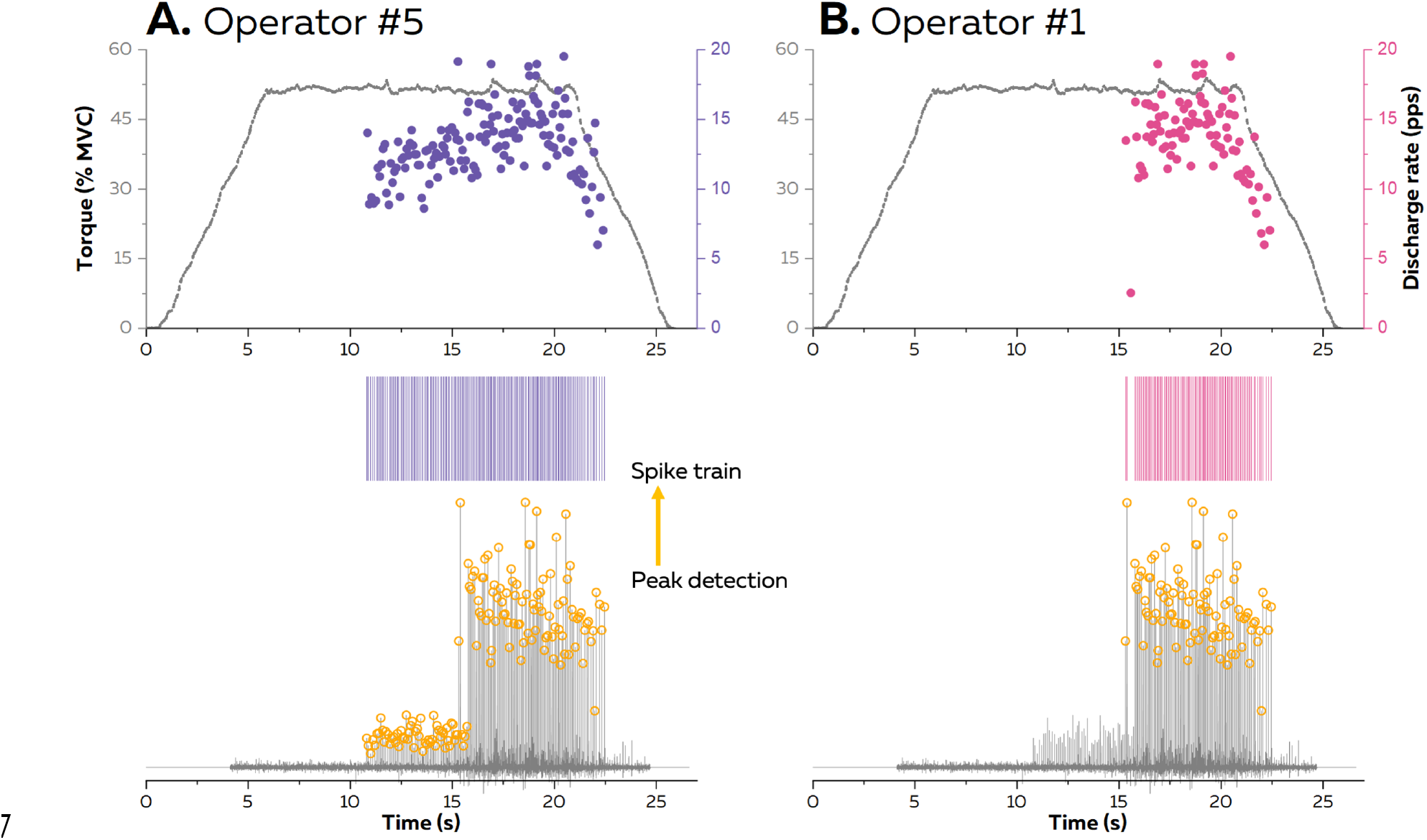
Example of discrepancies between two operators. The recalculation of the motor unit filter led to the identification of new spikes with moderate quality, which were selected by operator #5 but not by operator #1.

Both the RoA (99.0 ± 2.1 vs. 96.6 ± 5.5%; p < 0.001) and the pulse-to-noise ratio (35.7 ± 3.8 vs. 33.1 ± 3.6 dB; p<0.001) were higher for the motor units that were retained by all the 8 operators (n = 112) than those retained by only some operators (n = 24).

Fig. 5 shows the mean RoA value for each motor unit and each contraction. The association between the RoA and the PNR of the edited units was analyzed by correlation analysis. There was a significant positive correlation, albeit small (P = 0.026, r = 0.19), indicating that the inter-operator agreement tended to be higher for motor units that were identified with higher accuracy.

**Fig 5.**
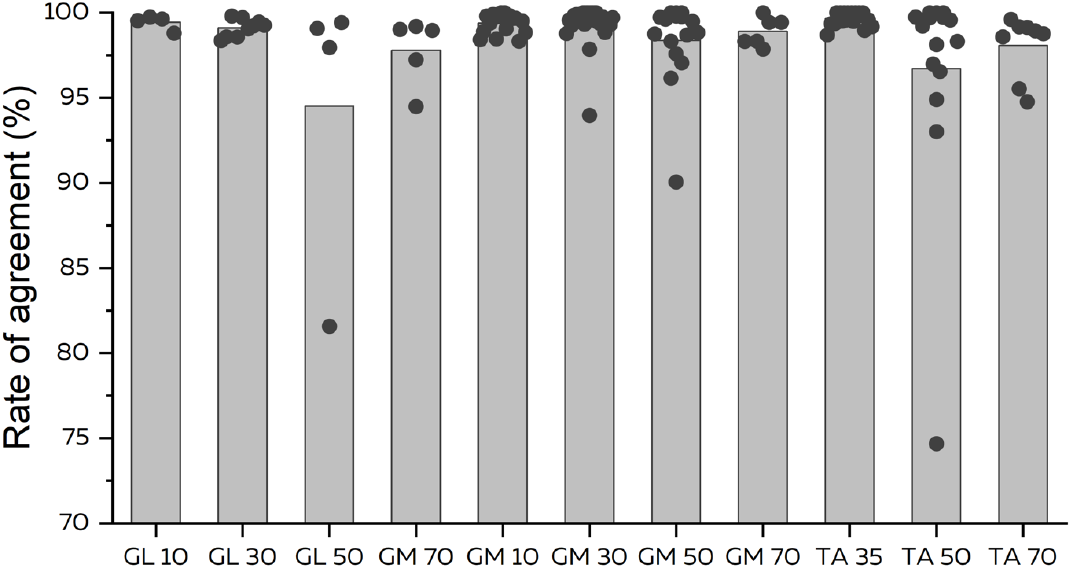
Rate of agreement between operators for each contraction and each motor unit. The mean rate of agreement across operators is depicted for each motor unit (black dot) and each contraction (grey bar). GL, Gastrocnemius lateralis; GM, Gastrocnemius medialis; TA, Tibialis anterior; Numbers indicate the contraction intensity in % of the maximal voluntary contraction.

We assessed the impact of the discrepancies between operators, albeit very limited, on two discharge characteristics commonly used, i.e. the mean discharge rate over the plateau and the time of recruitment. To do this, we considered the motor units identified by all the operators (n = 112). For both the mean discharge rate and the time of recruitment, the ICC value was very high (> 0.99).

## Discussion

We assessed the inter-operator reliability of identification of motor unit spike trains from HDsEMG. Specifically, eight operators were provided with the same automatically extracted data and were asked to manually edit them. Based on the 126 ± 5 motor units retained after manual edition, the median RoA value was very high (99.6%) leading to an excellent inter-operator reliability of the mean discharge rate at the force/torque plateau and the time at recruitment. There was a significant, albeit weak, positive correlation between RoA and the accuracy of the final decomposition assessed by the pulse-to-noise ratio. Taken together, these results show that identification of motor unit discharge times from HDsEMG decomposition is highly reliable across operators with varying levels of experience.

Recent advances in technology and signal processing have made it possible to identify motor units from surface HDsEMG signals. There is a growing interest in using this approach to improve our knowledge on the neural control of movement and to develop human-machine interfaces. However, the reliability of this approach is sometimes questioned, especially because it requires manual editing of the output from the decomposition algorithm (Enoka, 2019). As described in the methods section, manual editing requires different subjective steps (Del Vecchio et al., 2020), each having the potential to introduce discrepancies between operators. In our experiment, the eight operators analyzed the same dataset (extraction data) by following the same instructions, such as discarding motor units with a pulse-to-noise ratio < 30 dB. Despite this standardized process, they retained a slightly different number of motor units. There are different potential explanations for the differing number of motor units identified among operators. First, some operators could have discarded some motor units early in the editing process, considering that manual editing would not increase their pulse-to-noise ratio, while other operators could have successfully edited these same motor units. Second, the subjective steps of the editing process may have differed among operators, even slightly, such that only some operators succeeded in obtaining a pulse-to-noise ratio > 30 dB for some motor units. Specifically, the motor unit filter is updated from the spikes being present in the time-window of interest. Therefore, even small discrepancies in the selection of the false and true positives affect the motor unit filter and thus the spikes that will be identified with the updated filter. As both the most experienced operator (#8) and one of the least experienced operator (#1) extracted a similar number of motor units (n = 130, Table 3), we believe that experience did not have a major influence on the number of identified motor units, at least in our experimental conditions. Of note, quantifying the experience of the operators was not straightforward and further work is needed to assess the effect of experience and/or training on the editing step.

The vast majority of motor units were identified by all the operators. An important result is that the RoA between pairs of operators was very high, with a median value of 99.6% (Fig. 3). This means that when provided with the same decomposed data and the same basic instructions, operators converged toward almost identical motor unit spike trains. This logically led to very similar mean discharge rate and time of recruitment (both ICC > 0.99), which are two motor unit discharge characteristics often calculated from HDsEMG decomposition. In the rare cases where RoA values were low (0.6% of the RoA values < 85%), there was either a mistake made by one operator (motor unit #15 for GM 50% of MVC) or clear discrepancies between operators for a significant portion of the signal (motor unit #4 for GL 50% of MVC; Fig. 4). Specifically, Fig. 4 shows that the recalculation of the motor unit filter yielded to pulse trains of low amplitude, which were selected as being spikes by two out of the eight operators. Of note, because our study was designed to test the reliability between operators, we cannot infer accuracy; this means that we cannot determine which operators made the correct choice.

It is well known that the outcomes of the decomposition (number of identified units and accuracy) depend on the muscle being investigated, properties of the subcutaneous layers, and the contraction intensity (Del Vecchio et al., 2020). Here, we selected data from different muscles, different submaximal contraction intensities, and different participants. The number of identified motor units tended to be lower for GL than either GM or TA. In addition, there were less identified motor units at 70-75% of MVC than at lower intensities. Because of the small number of identified units for some muscles/contraction intensities, and the different contraction intensities tested for TA compared to GM and GL, it was impossible to systematically test for between-muscle or between-intensity differences in RoA, with appropriate statistics. However, given the very high RoA values (Fig. 5), we can confidently conclude that the inter-operator reliability was high, regardless of the muscle or the contraction intensity.

This study requires consideration of several methodological aspects. First, the high RoA values should be interpreted within the context of our standardized procedure, in which we gave a set of instructions based on previous recommendations (Del Vecchio et al., 2020). Slight divergence from this procedure, such as a different pulse-to-noise ratio threshold, could have led to different results. Importantly, our results demonstrate that following our basic set of instructions (see Methods) ensures that the manual analysis of motor unit spike trains is highly reliable across operators. Second, even though our results are based on outputs from a specific decomposition algorithm (DEMUSE tool software, see Methods), we do not believe that different results would have been obtained with another decomposition algorithms. Third, we selected operators with varying experience. Quantifying this experience is not straightforward. Ideally, we should have used the total number of units they had decomposed in their careers, but none of the operators had kept such a record. We therefore estimated this experience based on the number of published peer-reviewed articles including HDsEMG decomposition. Also, we selected operators from different research teams/universities, which ensured that they had had different prior training. Because HDsEMG decomposition requires training and some basic knowledge on neuromuscular physiology, we chose not to select fully inexperienced operators. Our results showed that moderate experience together with simple instructions are sufficient to ensure high reliability across operators. It suggests that the level of experience is certainly not a critical issue, at least not more than what is required for editing intramuscular EMG decomposition. We hope that the raw and the decomposed data provided with this article can be used to train new operators. Fourth, our experiment only considers data collected in males. We were not avoiding studying females; but for unknown reasons automatic decomposition is often more challenging in females (Del Vecchio et al., 2020). However, as we tested a large number of motor units with varying accuracy (as assessed by the initial pulse-to-noise ratio), we believe that reliability of manual editing would have been similar for signals collected in females, if automatic decomposition was successful. Finally, it is important to note that the high RoA across operators does not provide information about the decomposition accuracy. However, the accuracy of decomposition was verified in previous studies using experimental (Chen and Zhou, 2016; Holobar et al., 2010; Negro et al., 2016) or simulated signals (Holobar et al., 2014; Holobar and Zazula, 2007). Together with the excellent reliability of the manual editing reported herein, it provides further evidence that motor unit spike trains can be reliably estimated from HDsEMG decomposition of recordings obtained during submaximal isometric contractions.

## Acknowledgements

François Hug is supported by a fellowship from the *Institut Universitaire de France* (IUF) and a travel grant from the *Société de Biomécanique*. Support was received from the French national research agency (ANR-19-CE17-002-01, COMMODE project; to FH), from Slovenian Research Agency (J2-1731, L7-9421 and P2-0041; to AH), and from the European Research Council Synergy Grant NaturalBionicS (contract #810346; to DF).

